# Assessing digital phenotyping to enhance genetic studies of human diseases

**DOI:** 10.1101/738856

**Authors:** Christopher DeBoever, Yosuke Tanigawa, Matthew Aguirre, Greg McInnes, Adam Lavertu, Manuel A. Rivas

**Affiliations:** Department of Biomedical Data Science, Stanford University, Stanford, CA, USA

## Abstract

Population-scale biobanks that combine genetic data and high-dimensional phenotyping for a large number of participants provide an exciting opportunity to perform genome-wide association studies (GWAS) to identify genetic variants associated with diverse quantitative traits and diseases. A major challenge for GWAS in population biobanks is ascertaining disease cases from heterogeneous data sources such as hospital records, digital questionnaire responses, or interviews. In this study, we use genetic parameters including genetic correlation to evaluate whether GWAS performed using cases in the UK Biobank ascertained from hospital records, questionnaire responses, and family history of diseases implicate similar disease genetics across a range of effect sizes. We find that hospital record and questionnaire GWAS largely identify similar genetic effects for many complex phenotypes and that combining together both phenotyping methods improves power to detect genetic associations. We also show that family GWAS using cases ascertained on family history of disease agrees with combined hospital record/questionnaire GWAS and that family history GWAS has better power to detect genetic associations for some phenotypes. Overall, this work demonstrates that digital phenotyping and unstructured phenotype data can be combined with structured data such as hospital records to identify cases for GWAS in biobanks and improve the ability of such studies to identify genetic associations.

## Introduction

Genome-wide association studies (GWAS) for binary phenotypes such as presence of a disease typically obtain cases via methods like recruitment through medical systems or archived medical samples and compare these cases to controls known to not have the disease or random population controls where the disease is present at its population prevalence^1^. However, recent studies have begun to rely on self-reported phenotypes collected via questionnaires and web or mobile phone applications^2–10^. Such "digital phenotyping" may be faster and cheaper than standard cohort study approaches, but the extent to which this approach agrees with more traditional phenotyping approaches for GWAS is largely unknown because previous attempts to estimate the agreement between the two phenotyping approaches have focused on a small number of top associations and have not systematically assessed agreement across the hundreds or thousands of variants likely associated with complex, polygenic traits. For instance, a genome-wide study of self-reported thrombosis events found strong agreement between the top associations displayed in Manhattan plots from their self-reported thrombosis GWAS compared to previous cohort-based studies^2^. Other studies have reported overlaps with genome-wide significant loci from cohort studies but have not investigated the extent to which genetic effects that did not reach genome-wide significance agree^11^.

In addition to self-reported phenotypes, GWAS have also been performed using family history of disease as a proxy for disease diagnosis^12,13^. This genome-wide association study by proxy (GWAX) approach can be useful for childhood or late onset diseases where participants are difficult to recruit and is particularly appealing for population biobanking efforts that include questionnaires that ask about family history of disease. However, the degree to which proxy phenotyping attenuates effect sizes relative to traditional GWAS and the statistical power benefits of using GWAX in biobanks has not been explored. Estimating the agreement between digital phenotyping, GWAX, and traditional GWAS is important for understanding the extent to which these new phenotyping strategies may help uncover the genetic basis of human diseases and empower the generation of therapeutic hypotheses by, for instance, identifying strong acting protein-truncating variants^14–19^.

To explore the extent to which digital phenotyping or GWAX and traditional phenotyping approaches capture similar disease genetics, we developed a novel model called the multivariate polygenic mixture model (MVPMM) that estimates genetic parameters such as genetic correlation, polygenicity, and scale of genetic effects and applied the model to GWAS summary statistics from phenotypes in the UK Biobank whose cases were defined using hospital records, questionnaire responses, or family history information. We applied MVPMM to GWAS summary statistics from 41 binary medical phenotypes and found that there is strong agreement between the two phenotyping methods for most complex phenotypes. We then explored the extent to which combining these two phenotyping methods improves statistical power for GWAS. We next used MVPMM to compare how well GWAX agrees with these combined case definitions for a subset of phenotypes and found that family history GWAS has better power to detect associations in the UK Biobank for chronic bronchitis/emphysema, diabetes, and Alzheimer’s disease. The results from our study demonstrate that digital phenotyping and GWAX are useful approaches for identifying cases in large biobanks and can provide increased power for identifying associations for many conditions.

## Results

### Phenotyping, GWAS, and genetic parameter estimation

In order to perform GWAS and estimate genetic parameters, we stratified 337,199 European ancestry UK Biobank subjects into cases and controls for 41 binary medical phenotypes using hospital records or verbal questionnaire responses available from the UK Biobank (Table S1)^20^. The hospital records consist of hospital in-patient records (National Health Service Hospital Episode Statistics), cancer diagnoses from national cancer registries, and cause of death from national death registries. The verbal questionnaire data consisted of a computer survey that asked participants whether they had a history of several different illnesses followed by a verbal interview with a nurse to gain further confirmation of the selected diagnosis. The number and total fraction of cases ascertained from hospital records or questionnaire responses differed between phenotypes though each phenotype had at least 500 cases ascertained from each method (Figure 1A-B, Table S1). More than 80% of cases were identified using only one of the phenotyping methods for 20 of the phenotypes while 10 phenotypes had substantial overlap (>33%) in cases identified by both hospital records and verbal questionnaire data. Overall, however, 32/41 and 20/41 phenotypes had at least 25% of cases derived solely from hospital records or questionnaire data, respectively, indicating that both phenotyping methods add a substantial proportion of cases for most diseases (Figure 1A-B).

**Figure 1.**
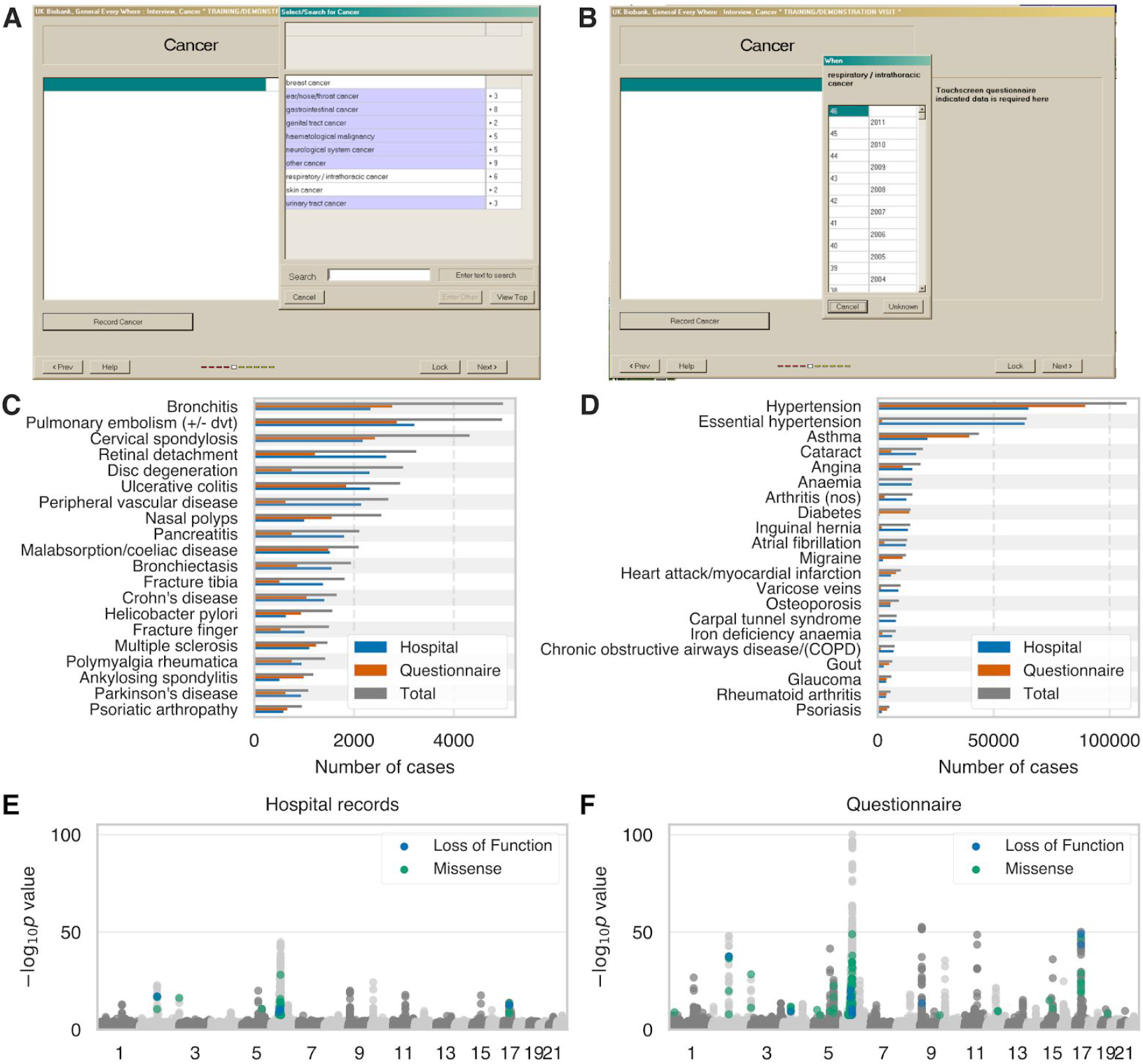
(A,B) Screenshot of UK Biobank questionnaire where participants can indicate that they have been diagnosed with specific (A) cancers or other illnesses and (B) specify at what age they were diagnosed. (C,D) Number of cases for each of 41 medical phenotypes where cases are defined using hospital records (blue), questionnaire responses (orange), or both combined (grey). (E,F) Manhattan plots for asthma GWAS with cases defined by (E) hospital records or (F) questionnaire responses. Loss of function and missense variants with p<5e-8 are colored blue and green, respectively. Grey dots indicate all other variants.

For each phenotype, we used cases defined by either hospital records or verbal questionnaire responses to perform GWAS for 784,257 variants genotyped by array (Methods). For example, for asthma, we defined 21,445 cases using hospital records and 39,483 cases using questionnaire responses, of which 17,302 cases were shared between both phenotyping methods. Performing GWAS for each phenotyping method yielded similar Manhattan plots, though there is less power to detect associations for hospital records as expected due to the lower number of cases (Figure 1C-D). For instance, the p-value for the reported association between the protein truncating variant rs146597587 in *IL33* and asthma is 7.1×10^−7^ for hospital records and 2.4×10^−14^ for questionnaire responses^21^. While these results illustrate the usefulness of the verbal questionnaire data, it is difficult to draw conclusions about the overall agreement between the two phenotyping methods outside of the small number of top GWAS findings.

To estimate the agreement between phenotyping using hospital records and verbal questionnaire responses for identifying genetic associations, we applied a novel Bayesian mixture model, the MultiVariate Polygenic Mixture Model (MVPMM), to GWAS summary statistics (effect size estimate and standard error of effect size estimate) for the 41 medical phenotypes where cases were defined for each phenotype using either hospital in-patient records or self-reported verbal questionnaire responses (Table S1). MVPMM estimates genetic parameters including genetic correlation, polygenicity, and scale of effect sizes by modeling GWAS summary statistics as drawn from either a null component where the true effect of the variant on the phenotype is zero or a non-null component where the true effect of the variant on the phenotype is non-zero. For both components, summary statistics (treated as the data) for each variant are modeled as being drawn from a multivariate normal distribution with zero mean and unknown covariance matrix. For the null component, the covariance matrix uses the standard error of the effect size estimate and estimates the correlation of errors that may be due to shared subjects. The covariance matrix for the non-null component combines the error covariance matrix from the null component with another covariance matrix that captures the genetic correlation between the phenotypes being considered. This model allows us to estimate the (1) genetic correlation between two phenotypes, (2) fraction of loci that belong to the non-null component for both phenotypes (polygenicity), and (3) scale of the genetic effects for each phenotype (Methods).

### GWAS based on hospital records or questionnaire responses

To systematically examine whether the GWAS results for phenotyping using hospital records or questionnaire responses agreed across a broader range of associations, we applied MVPMM to the GWAS summary statistics for the 41 phenotypes to estimate the genetic correlation between the results of both phenotyping methods (Figure 2, Table S2, Methods). The genetic correlation estimates from MVPMM were robust according to the 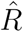 statistic (Figure S1) and agreed in large part with those from LD score regression (Figure S2)^22^. We found that 21/41 phenotypes had genetic correlations greater than 0.9 indicating strong agreement of genetic effects between cases identified by hospital records or verbal questionnaire data. Another 6/41 phenotypes had genetic correlations greater than 0.8 indicating moderate agreement between the two phenotyping methods. For instance, the genetic correlation between asthma as defined by the phenotyping methods was 0.96 (95% highest posterior density (HPD) 0.95-0.98, Methods). We identified 43,626 total asthma cases between both phenotyping methods, 40% of which were identified by both methods and 51% of which were identified only by the verbal questionnaire responses. These results indicate that the large number of asthma cases contributed by the verbal questionnaire responses capture similar disease genetics as the cases indicated by hospital records. We observed similar results for several other diseases where a large number of cases were identified from questionnaire responses such as ankylosing spondylitis, psoriasis, myocardial infarction, gout, and others (Figure 2) demonstrating that the two phenotyping methods agree for a range of phenotypes including both chronic and acute conditions.

**Figure 2.**
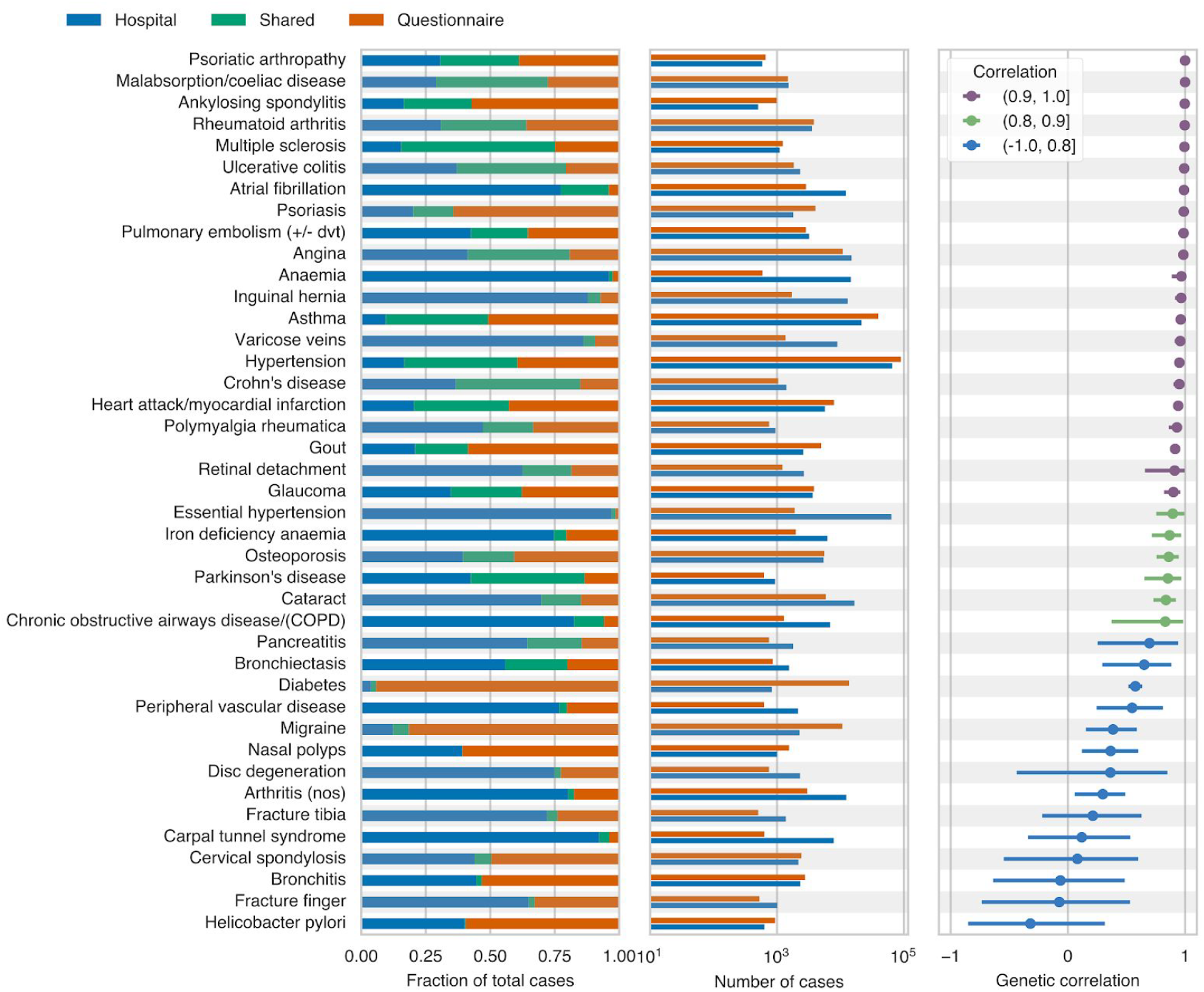
The first panel from left indicates the fraction of cases that were ascertained from hospital records only (blue), questionnaire responses only (orange), or both phenotyping methods (green). The second panel shows the number of cases ascertained from hospital records (blue) and questionnaire responses (orange). The third panel shows the estimated genetic correlation from MVPMM; the dot shows the mean of the posterior distribution and the bars show the 95% highest posterior density (Methods).

There were 14 phenotypes had genetic correlations less than 0.8 indicating less agreement between cases defined by hospital records or questionnaire data, though notably several of these pairs were predicted to have positive, non-zero correlations (Figure 2). For instance, the genetic correlations for migraine (0.38, 95% HPD: 0.15-0.59), peripheral vascular disease (0.55, 95% HPD: 0.25-0.81), and carpal tunnel syndrome (0.19, 95% HPD: −0.34-0.53) were all less than 0.8 indicating that there may be differences in the case populations captured by the phenotyping methods for these diseases. The Manhattan plots for these phenotypes are also different for the two phenotyping methods demonstrating that even the top associations are not necessarily consistent between the two methods for these phenotypes (Figure S3). These results indicate that there may be some differences in the case populations captured by hospital records and questionnaire responses for these phenotypes.

Given the high genetic correlation between the two phenotyping methods for many of the phenotypes tested here, we combined together cases from both phenotyping methods, performed GWAS analysis using the combined cases, and used MVPMM to estimate genetic parameters between GWAS summary statistics from combined cases and questionnaire cases or hospital record cases (Table S2). We found a high correlation between the combined GWAS and GWAS using either questionnaire cases or hospital record cases. 29 phenotypes had genetic correlations greater than 0.8 for the hospital record GWAS, and 33 phenotypes had correlations greater than 0.8 for the verbal questionnaire GWAS. We compared the estimates from MVPMM for the scale of effects, which captures how strong the genetic effects are for each phenotype definition, and found that the scale of effects generally agreed between the combined GWAS and the questionnaire or hospital record GWAS (Figure 3A-B) indicating that there is not a large amount of effect size attenuation due to combining the phenotyping methods. We calculated the power to detect associations using the combined cases compared to either the questionnaire or hospital record cases and found an increase in power for detecting associations for both risk and protective rare variants (Figure 3C-D, Figures S4–S5). The increase in power differs across phenotypes depending on the fraction of total cases that are added by including cases ascertained from questionnaire data. Notably, identifying additional cases causes a larger increase in the power to detect rare protective variants which are especially useful for identifying therapeutic targets^14,17,18,23^.

**Figure 3.**
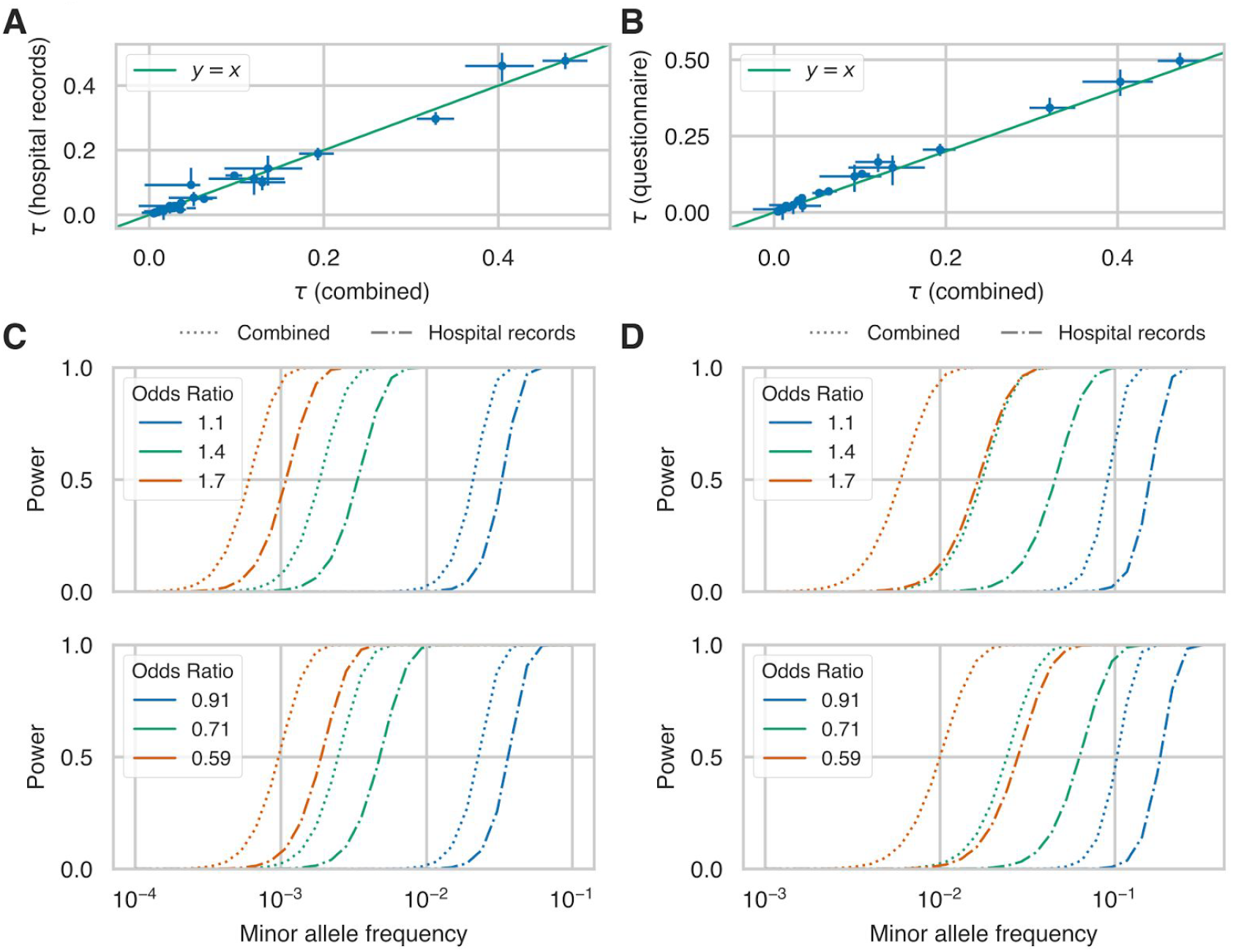
(A,B) Estimates of scale of genetic effects (*τ*) from MVPMM for GWAS summary statistics generated using cases ascertained from hospital records and verbal questionnaire (combined) versus summary statistics generated using only (A) hospital record or (B) verbal questionnaire cases. Phenotypes whose 95% HPD size for *τ* was less than 0.1 for both hospital record and verbal questionnaire comparisons are plotted. (C,D) Statistical power to detect association between rare genetic variants at different minor allele frequencies for (C) asthma and (D) psoriasis in the UK Biobank. Dot-dash lines show power for GWAS performed using only cases ascertained from hospital records and dotted lines show power for GWAS performed using cases ascertained from both hospital records and verbal questionnaire data. Top panel shows power for rare risk variants and bottom panel shows power for rare protective variants. Different colors indicate power for different association effect sizes.

### GWAS using disease diagnosis or family history of disease

Another approach for identifying loci associated with disease is a genome-wide association study by proxy (GWAX) where cases are defined as biobank participants that have a relative with a particular disease^12,13^. We estimated genetic parameters for 15 diseases using summary statistics from a traditional GWAS where cases were identified either from hospital records and/or questionnaire responses and summary statistics for the same disease from a GWAX based on the presence of disease in the parents of the subject (ascertained from questionnaire data). We included multiple disease definitions for diabetes and emphysema that rely on different aspects of the UK Biobank phenotyping data. We restricted our analysis to diseases with at least 1,000 GWAS cases (except for Alzheimer’s disease), though notably, the number of cases is generally much larger for GWAX than GWAS. We found that the genetic correlation was greater than 0.9 for 10/15 comparisons while four comparisons had genetic correlations less than 0.8. One of the comparisons with genetic correlation less than 0.8 was family history of “severe depression” and mania/bipolar disorder/manic depression. In this case, these two case definitions were matched due to the word “depression” but actually capture two different diseases, depression and bipolar disorder, and the low genetic correlation reflects this. Another comparison with genetic correlation less than 0.8 is type 1 diabetes and family history of diabetes. However, family history of diabetes has a high correlation with other diabetes definitions that likely include type 2 diabetes cases, indicating that family history of diabetes mostly captures cases for type 2 diabetes, consistent with the higher prevalence of type 2 diabetes in the UK Biobank^24^.

Since our GWAX uses subjects whose parents had a particular disease, we expect that the effect sizes of the associated variants identified by GWAX will be attenuated relative to the effect sizes estimated from GWAS^13^. We used the scale of effects estimates for each phenotype to estimate the attenuation for GWAX compared to combined hospital record/verbal questionnaire GWAS for the 10 phenotypes with genetic correlations greater than 0.9. We found that the estimated attenuation factors ranged from 0.24-0.54 and that observed effect sizes were generally scaled consistent with the estimated attenuation factor (Figure 5A-C, Figure S6). While the smaller effect sizes of GWAX may decrease the power to detect genetic associations compared to GWAS in the UK Biobank, we find that this decrease in power is offset by much larger case sizes in GWAX for some phenotypes (Figure 5D-E, Figures S7–S8). For instance, the power to detect associations for chronic bronchitis/emphysema, diabetes, and Alzheimer’s disease is higher using GWAX whereas the power to detect associations is higher for combined hospital record/verbal questionnaire GWAS for other phenotypes, such as prostate cancer.

**Figure 4.**
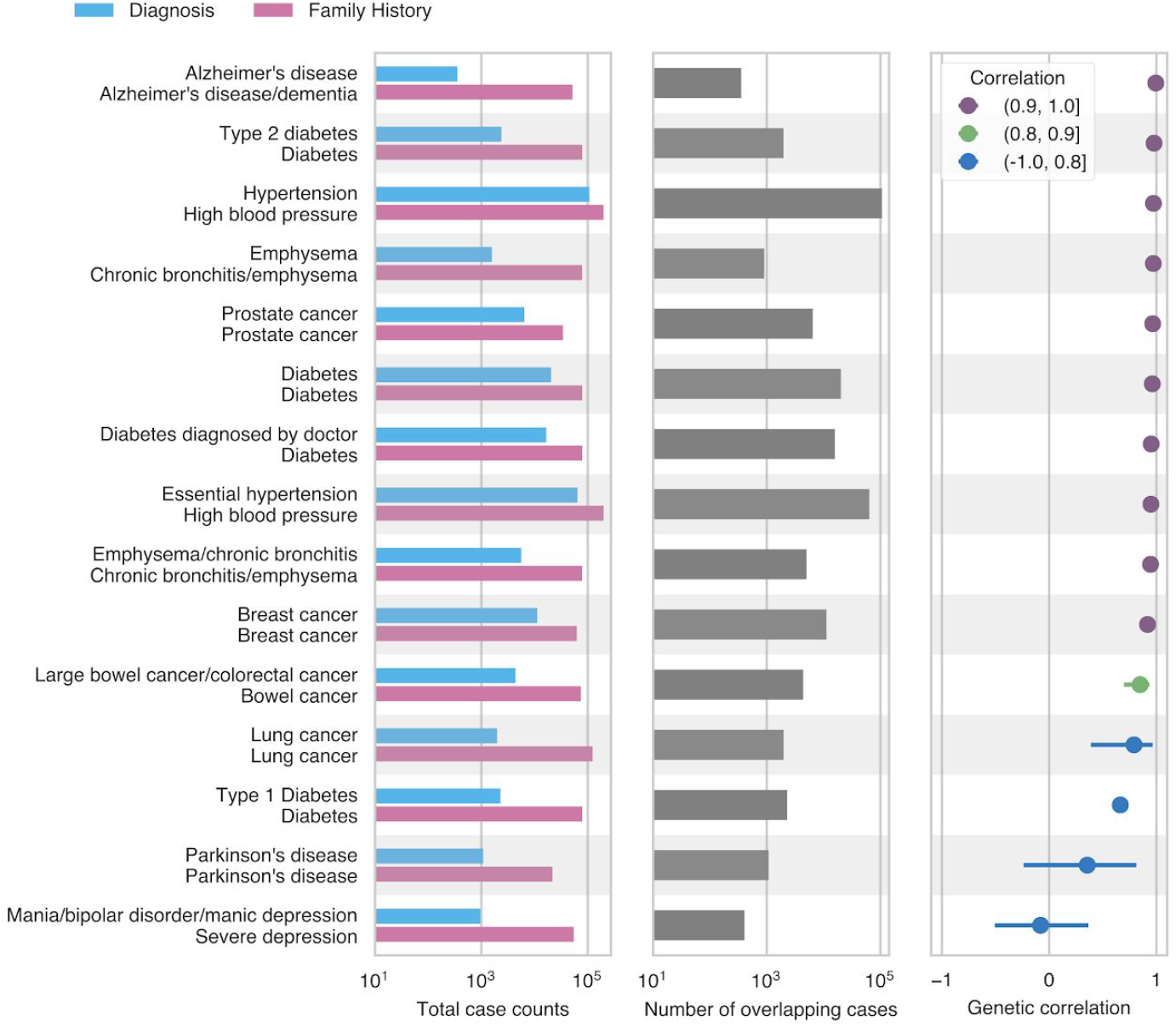
The first panel from left indicates the number of cases ascertained from combined hospital records and questionnaire responses (blue) or family history of disease (pink). The second panel shows the number of cases overlapping between both phenotyping methods. The third panel shows the estimated genetic correlation between GWAS and GWAX from MVPMM; the dot shows the mean of the posterior distribution and the bars show the 95% highest posterior density (Methods).

**Figure 5.**
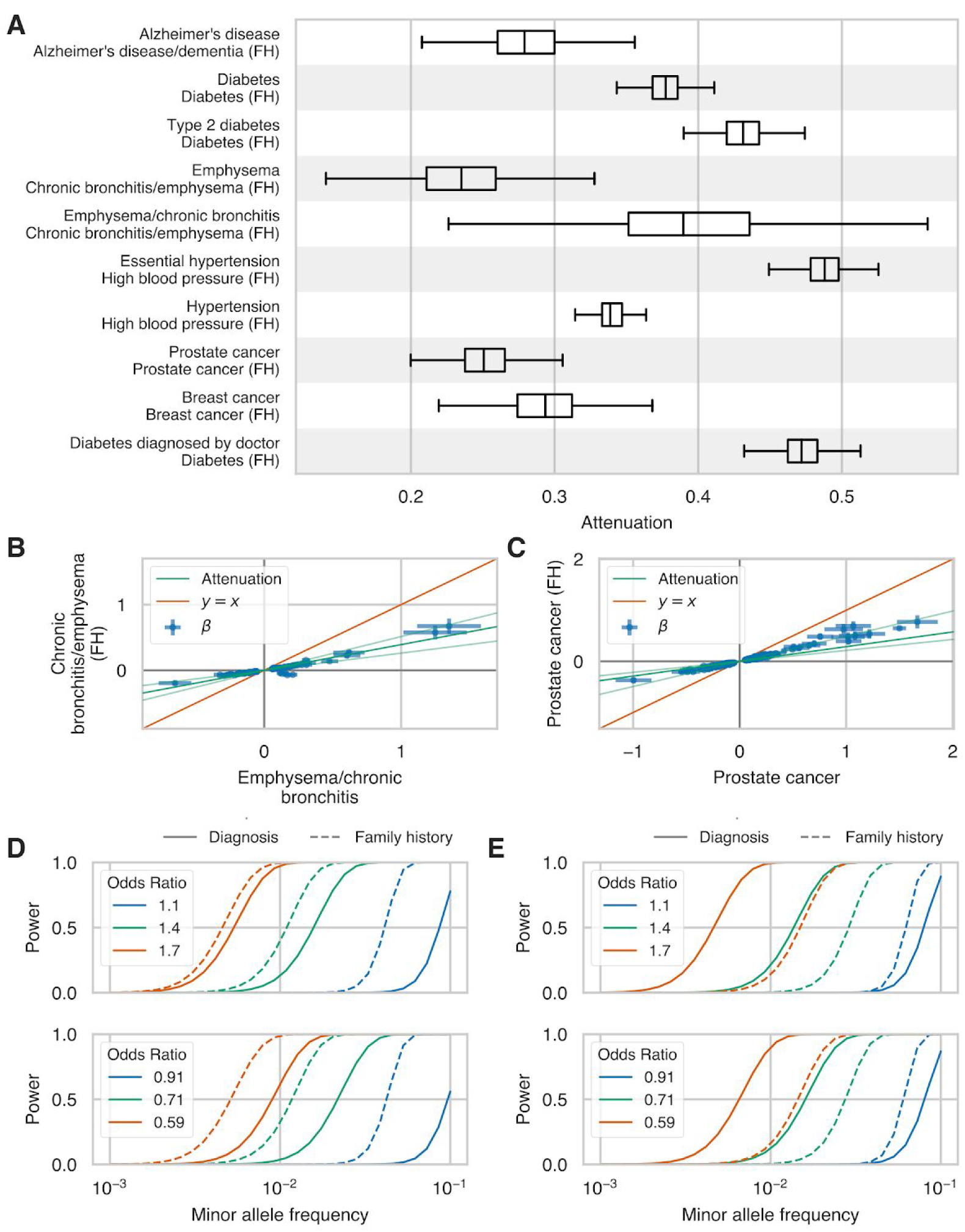
(A) Box plots of the posterior distribution of effect size attenuation (Methods) from GWAX using cases ascertained on family history of disease versus GWAS using cases ascertained using combined hospital records and verbal questionnaire responses. (B, C) Effect sizes (*β*) and standard errors for family history GWAX (y-axis) versus combined hospital records and verbal questionnaire responses GWAS (x-axis) for (B) chronic bronchitis/emphysema and (C) prostate cancer. Dark green line indicates the mean of the posterior distribution of attenuation from MVPMM and light green lines indicate lower and upper bounds of 95% highest posterior density of attenuation (Methods). (D, E) Statistical power to detect association between rare genetic variants at different minor allele frequencies for (D) chronic bronchitis/emphysema and (E) prostate cancer in the UK Biobank. Solid lines show power for GWAS performed using cases ascertained from hospital records and questionnaire responses and dashed lines show power for GWAS performed using cases ascertained from using or family history of disease. Top panel shows power for rare risk variants and bottom panel shows power for rare protective variants. Different colors indicate power for different association effect sizes.

## Discussion

In this study, we present a novel method for estimating genetic parameters from GWAS summary statistics called the MultiVariate Polygenic Mixture Model (MPVMM) and use the method to evaluate the extent to which GWAS using cases ascertained from hospital records, verbal questionnaire responses, and family history of disease agree across 41 diverse medical phenotypes. We found that GWAS using cases ascertained from hospital records or questionnaire responses had genetic correlation greater than 0.8 for 27 phenotypes indicating that the two phenotyping methods identify similar disease genetics for complex diseases.

Combining both phenotyping methods for GWAS does not greatly alter effect size estimates relative to using either method individually but does increase power to identify genetic associations due to the increased number of cases. We also showed that GWAX, where family history of disease is used to identify cases, has genetic correlation greater than 0.8 with combined hospital record/questionnaire GWAS for 11 of 16 pairs of traits analyzed demonstrating the GWAX approaches based on digital phenotyping can also be used to identify variant-disease associations. Finally, we showed that the power to detect genetic associations in the UK Biobank is greater for GWAX than GWAS for chronic bronchitis/emphysema, diabetes, and Alzheimer’s disease. The high genetic correlation between GWAS based on questionnaire data and GWAS based on hospital records shows that the two methods capture similar disease genetics. In the UK Biobank, participants completed a touchscreen questionnaire and had a follow-up interview with a nurse to discuss any diagnoses for major illnesses and procedures. Future studies will explore to what extent other digital or questionnaire phenotyping approaches such as phone or internet applications, waiting room surveys, or features extracted by natural language processing also identify similar disease genetics to GWAS that ascertain cases using more traditional recruitment methods. Such comparisons may be aided by the adoption of standardized questionnaire approaches across datasets or biobanks so digital phenotyping methods can easily be shared in the way that phenotyping based on structured medical data are now shared^25^. The results from this study illustrate how such efforts will benefit GWAS in population-scale biobanks by improving power to detect novel genetic associations.

## Author Contributions

M.A.R. conceived and designed the study. C.D., Y.T., M.A., G.M., A.L., and M.A.R. designed and carried out the statistical and computational analyses. C.D., M.A., Y.T., G.M., and A.L. carried out quality control of the data. The manuscript was written by C.D. and M.A.R. M.A.R. supervised all aspects of the study.

## Acknowledgments

This research has been conducted using the UK Biobank resource. We thank all the participants in the UK Biobank study. M.A.R. is supported by Stanford University and a National Institute of Health center for Multi- and Trans-ethnic Mapping of Mendelian and Complex Diseases grant (5U01 HG009080). C.D. is supported by a postdoctoral fellowship from the Stanford Center for Computational, Evolutionary, and Human Genomics and a seed grant from the Stanford ChEM-H Institute. Y.T. is supported by Funai Overseas Scholarship from Funai Foundation for Information Technology and the Stanford University Biomedical Informatics Training Program. The primary and processed data used to generate the analyses presented here are available in the UK Biobank access management system (https://amsportal.ukbiobank.ac.uk/) for application 24983, “Generating effective therapeutic hypotheses from genomic and hospital linkage data” (http://www.ukbiobank.ac.uk/wp-content/uploads/2017/06/24983-Dr-Manuel-Rivas.pdf), and the results are displayed in the Global Biobank Engine (https://biobankengine.stanford.edu). Research reported in this publication was supported by the National Human Genome Research Institute of the National Institutes of Health under Award Number R01HG010140. The content is solely the responsibility of the authors and does not necessarily represent the official views of the National Institutes of Health

## Methods

### Quality Control of Genotype Data

We used genotype data from UK Biobank data set release version 2 for all aspects of the study^26^. To minimize the impact of cofounders and unreliable observations, we used a subset of 337,199 individuals that satisfied all of the following criteria: (1) self-reported white British ancestry, (2) used to compute principal components, (3) not marked as outliers for heterozygosity and missing rates, (4) do not show putative sex chromosome aneuploidy, and (5) have at most 10 putative third-degree relatives. We used PLINK v1.90b4.4^27^ to compute the following statistics for each of 784,257 variants: (a) genotyping missingness rate, (b) p-values of Hardy-Weinberg test, and (c) allele frequencies. As described previously^14^, we removed variants that had (1) missingness rate greater than 1%, (2) Hardy-Weinberg disequilibrium test p-value less than 1e-7, (3) ambiguous cluster plots, or (4) minor allele frequencies inconsistent with gnomAD.

### Hospital Record and Verbal Questionnaire Phenotype Definitions

We used the following procedure to define cases and controls for non-cancer phenotypes. For a given phenotype, ICD-10 codes (Data-Field 41202) were grouped with self-reported non-cancer illness codes from verbal questionnaires (Data-Field 20002) that were closely related. This was done by first creating a computationally generated candidate list of closely related ICD-10 codes and self-reported non-cancer illness codes, then manually curating the matches. The computational mapping was performed by calculating the token set ratio between the ICD-10 code description and the self-reported illness code description using the FuzzyWuzzy python package. The high scoring ICD-10 matches for each self-reported illness were then manually curated to ensure high confidence mappings. Manual curation was required to validate the matches because fuzzy string matching may return words that are similar in spelling but not in meaning. For example, to create a hypertension cohort the code description from Data-Field 20002 ("Hypertension") was mapped to all ICD-10 code descriptions and all closely related codes were returned ("I10: Essential (primary) hypertension" and "I95: Hypotension"). After manual curation code I10 would be kept and code I95 would be discarded. After matching ICD-10 codes and with self-reported illness codes, cases were identified for each phenotype using only the associated ICD-10 codes, only the associated self-reported illness codes, or both the associated ICD-10 codes and self-reported illness codes.

Questionnaire images were from the UK Biobank website at http://biobank.ctsu.ox.ac.uk/crystal/crystal/images/vs_when_1.png and http://biobank.ctsu.ox.ac.uk/crystal/crystal/images/vs_review_2.png.

### Family History Phenotype Definitions

We used data from Category 100034 (Family history - Touchscreen - UK Biobank Assessment Centre) to define "cases" and controls for family history phenotypes. This category contains data from the touchscreen questionnaire on questions related to family size, sibling order, family medical history (of parents and siblings), and age of parents (age of death if died). We focused on Data Coding 20107: Illness of father and 20110: Illness of mother.

### Cancer Phenotype Definitions

We combined cancer diagnoses from the UK Cancer Register with self-reported diagnoses from the UK Biobank questionnaire to define cases and controls for cancer GWAS. Individual level ICD-10 codes from the UK Cancer Register (http://biobank.ctsu.ox.ac.uk/crystal/label.cgi?id=100092), Data-Field 40006 (http://biobank.ctsu.ox.ac.uk/crystal/field.cgi?id=40006), and the National Health Service (http://biobank.ctsu.ox.ac.uk/crystal/label.cgi?id=2022), Data-Field 41202 (http://biobank.ctsu.ox.ac.uk/crystal/field.cgi?id=41202), in the UK Biobank were mapped to the self-reported cancer codes, Data-Field 20001 (http://biobank.ctsu.ox.ac.uk/crystal/field.cgi?id=20001). The mapping was performed via manual curation of ICD-10 codes for each of the self-reported cancer codes. UKB field codes for self-reported cancer were created with a tree structure such that more specific cancer subtypes (e.g. “malignant melanoma”) are nested under more general categories (“skin cancer”). This tree structure was preserved in the field code to ICD-10 mapping. For example, the self-reported phenotype of “lip cancer” was mapped to its field code, 1010, and the ICD-10 codes for “malignant neoplasm of lip”, C00 and C000-C009. After this mapping, individuals with an affirmative entry in one or more of the phenotype collections (self-reported cancer, cancer registry, and the NHS) were included in the case cohort for the GWAS. No secondary neoplasms were included in the cancer phenotype mappings.

### Genome-Wide Association Analyses

We performed genome-wide association analyses for binary medical phenotypes in the UK Biobank across 784,257 variants genotyped by array using logistic regression with Firth-fallback as implemented in PLINK v2.00a (17 July 2017). Firth-fallback is a hybrid algorithm which normally uses the logistic regression code described in (Hill 2017), but switches to a port of logistf() (https://cran.r-project.org/web/packages/logistf/index.html) in two cases: (1) one of the cells in the 2×2 allele count by case/control status contingency table is empty (2) logistic regression was attempted since all the contingency table cells were nonzero, but it failed to converge within the usual number of steps. We used the following covariates in our analysis: age, sex, array type, and the first four principal components, where array type is a binary variable that represents whether an individual was genotyped with UK Biobank Axiom Array or UK BiLEVE Axiom Array. For variants that were specific to one array, we did not use array as a covariate.

### Multivariate Polygenic Mixture Model (MVPMM)

#### Model Definition

We developed a two-component mixture model to estimate genetic parameters including correlation, scale, and proportion of non-zero genetic effects. Let 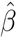 be an *N* × 2 matrix of estimated GWAS effect sizes (regression coefficient for quantitative phenotypes, log odds ratio for binary phenotypes) and let 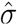 be an *N* × 2 matrix of estimated standard errors for *N* LD-independent loci for two phenotypes. Let *β*_*i*_ be a column vector with the effect sizes for the *i*th locus and let *σ*_*i*_ be a column vector with standard errors for the *i*th locus. Under the MultiVariate Polygenic Mixture Model (MVPMM), the observed effect sizes and standard errors are assumed to be generated from one of two mixture components. The first component is a point-mass at zero such that 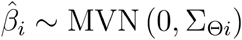 where 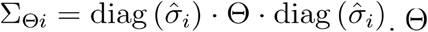 is a 2 × 2 correlation matrix describing the correlation between the observed effect sizes (measurement errors) at the null variants and 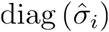 is a diagonal matrix with 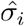 on the diagonal. The second component is a multivariate normal distribution with mean zero and unknown covariance matrix such that 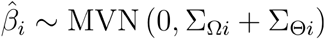 where Σ_Ω*i*_ = diag(*τ*) ⋅ Ω ⋅ diag(*τ*). Ω is a 2 × 2 correlation matrix describing the correlation between the genetic effects of the two phenotypes and *τ* is a length-2 vector describing the scale of the genetic effects. The model includes a mixing parameter *π* that describes the fraction of variants in the second component that captures variants associated with the two phenotypes. The LKJ prior was used for the 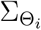 and 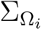 correlation matrices. The other priors are *τ*_*i*_ ~ cauchy (0,2.5), *π* ~ Dir (1).

#### Estimating Genetic Parameters using MVPMM

We implemented MVPMM using the Stan probabilistic programming language and used MVPMM to estimate genetic parameters for a given pair of GWAS summary statistics as follows. First, we obtained GWAS summary statistics (effect size and standard error) for 361,436 LD-independent autosomal variants; these 361,436 LD-independent variants were identified using plink (--indep 50 5 2). These variants were filtered to include only those whose standard error was less than 0.2 in both phenotypes to remove variants with uncertain effect size estimates. We then performed MCMC sampling using Stan with four chains for 500 iterations with 100 burn-in iterations. We calculated the 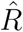 statistic for the genetic correlation parameter Ω_21_ to evaluate whether the MCMC sampling converged. If 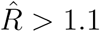, we repeated MCMC sampling with four chains for 1,000 iterations with 200 burn-in iterations. We excluded bronchiectasis for the combined phenotyping versus hospital record phenotyping (Figure 3A) because MCMC sampling was extremely slow (500 iterations did not finish in seven days). We ran MVPMM for GWAS using cases defined by either hospital records or questionnaire responses for 51 medical phenotypes. We then filtered out ten phenotypes with unrealistic polygenicity estimates (*π* > 0.4) that indicated model failure or which had 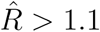 indicating that the MCMC sampling did not converge.

Parameter estimates are plotted as dots that indicate the mean of the posterior distribution and bars that show the 95% highest posterior density (HPD). The 95% HPD is the smallest interval that includes 95% of the density of the posterior distribution.

#### Effect Size Attenuation Estimates

We calculated the attenuation for each GWAX as 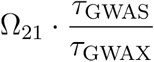 where Ω_21_ is the estimated genetic correlation between the GWAS and GWAX, *τ*_GWAS_ is the estimated scale parameter for the GWAS, and *τ*_GWAX_ is the estimated scale parameter for the GWAX. The attenuation was calculated for each MCMC sample to obtain a posterior distribution of the attenuation for each GWAS/GWAX pair.

For attenuation scatter plots, GWAS summary statistics and p-values were obtained for each GWAX/GWAS pair for the variants used as input to MVPMM. The following procedure was used to identify variants with reasonable effect size estimates to plot to demonstrate attenuation. Variants were filtered to include only those with *p* < 0.001 in both GWAS and GWAX. If there were less than 500 variants with *p* < 0.001 in both GWAS and GWAX, the p-value filter threshold was increased by a factor of two until there were at least 500 variants or the threshold exceeded 0.01. This resulted in a set of variants with effect sizes that could be compared between GWAX and GWAS.

### Power Calculations

Power calculations were performed using a forked version of the GeneticsDesign Bioconductor package (https://bioconductor.org/packages/devel/bioc/html/GeneticsDesign.html) available at https://github.com/cdeboever3/GeneticsDesign that adds support for unselected controls (i.e. disease status is present at population prevalence in controls) in case/control studies. Disease prevalence was estimated as the total number of cases identified by combining hospital records and verbal questionnaire responses and dividing by the total number of subjects. We calculated power curves for medical phenotypes with genetic correlation greater than 0.8 and GWAX phenotypes with genetic correlation greater than 0.9.

## Supplementary Materials

### Supplemental Figures

**Figure S1.**
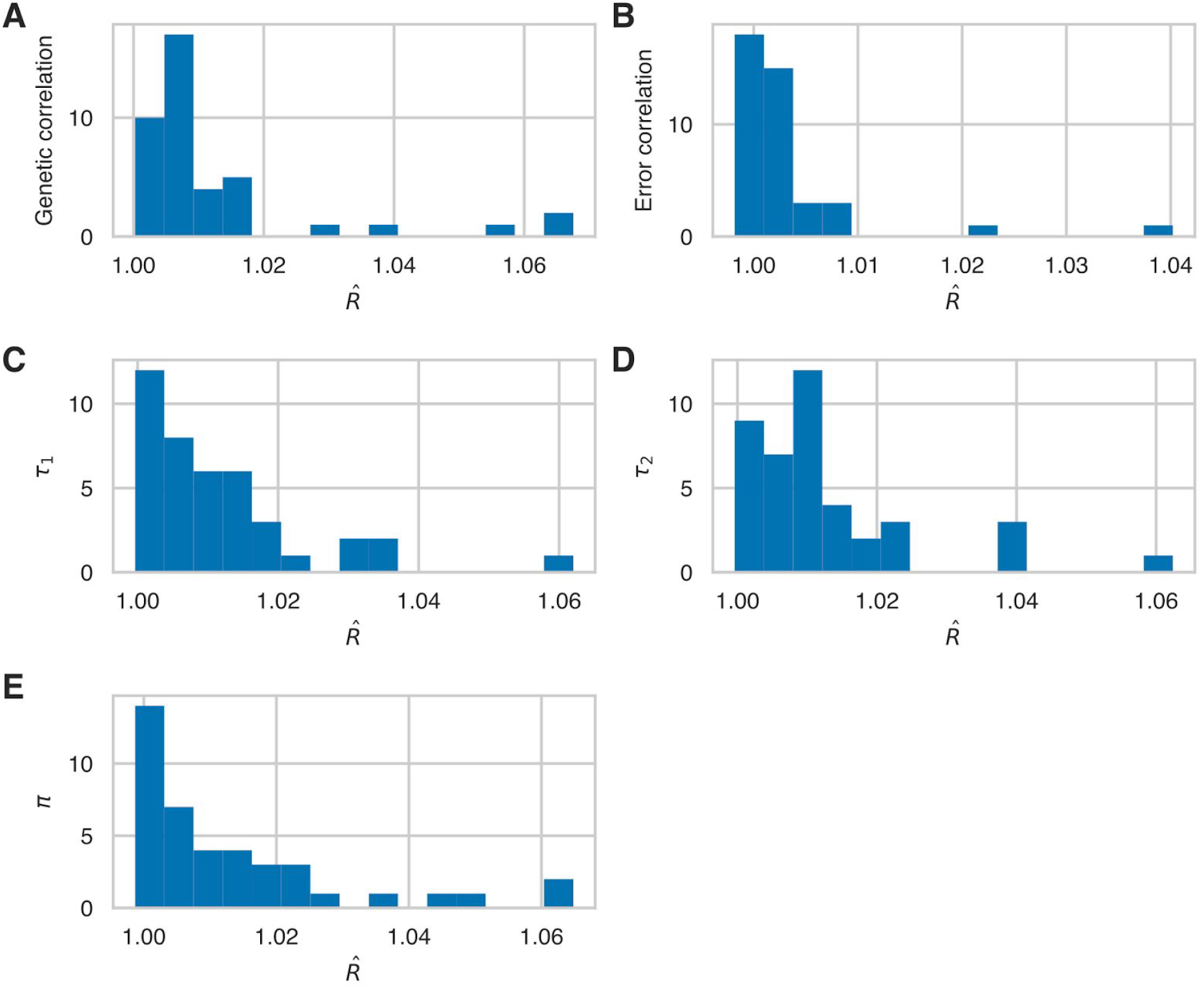
Histogram of 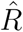 values demonstrating MCMC convergence for parameters estimated by MVPMM using GWAS summary statistics for 41 phenotypes where cases were defined using hospital records or verbal questionnaire responses.

**Figure S2.**
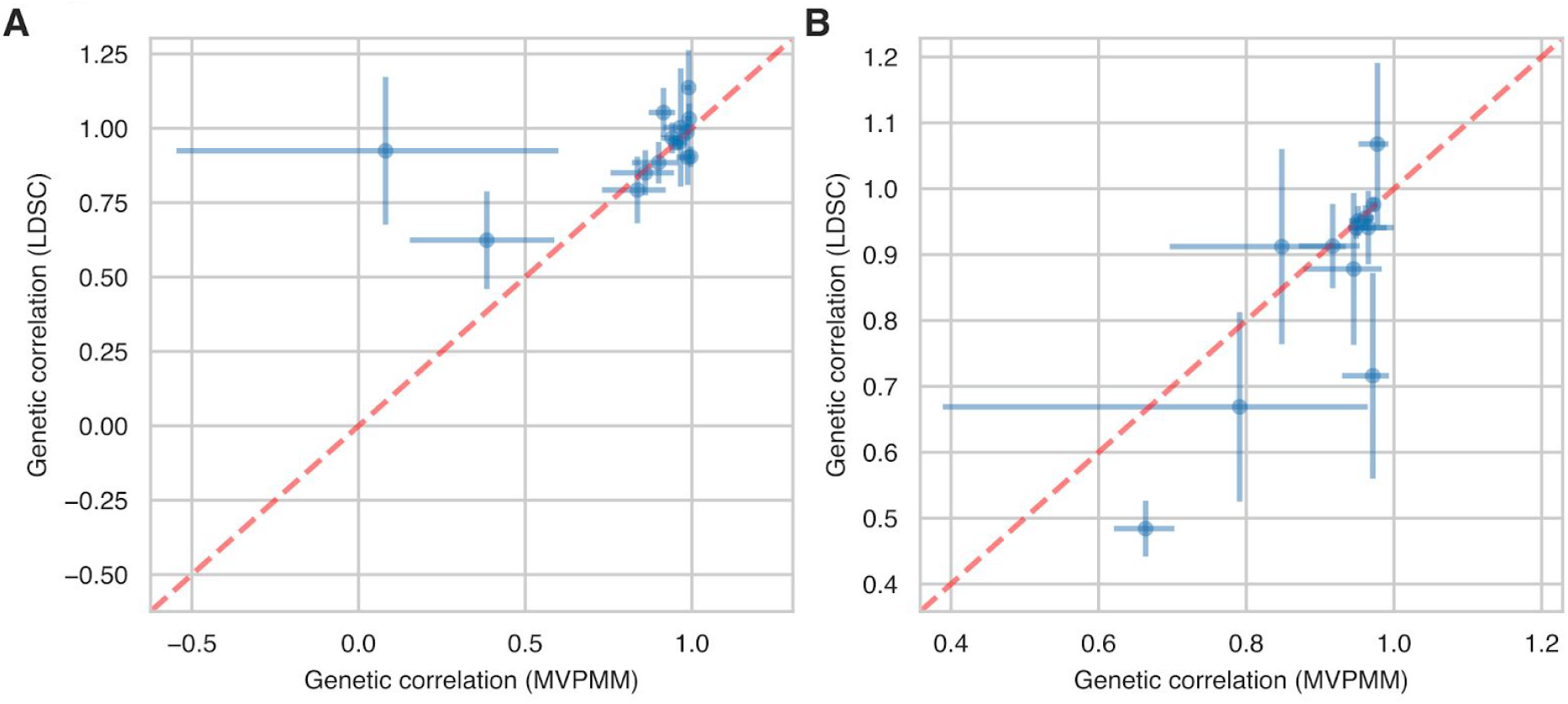
(A,B) Genetic correlation estimates from MVPMM (x-axis) and LD score regression (y-axis) using (A) GWAS summary statistics generated using disease definitions from hospital records or verbal questionnaire responses (minimum 1,500 cases for each) or (B) GWAS summary statistics from disease diagnosis or family history of disease (minimum 1,500 cases for each).

**Figure S3.**
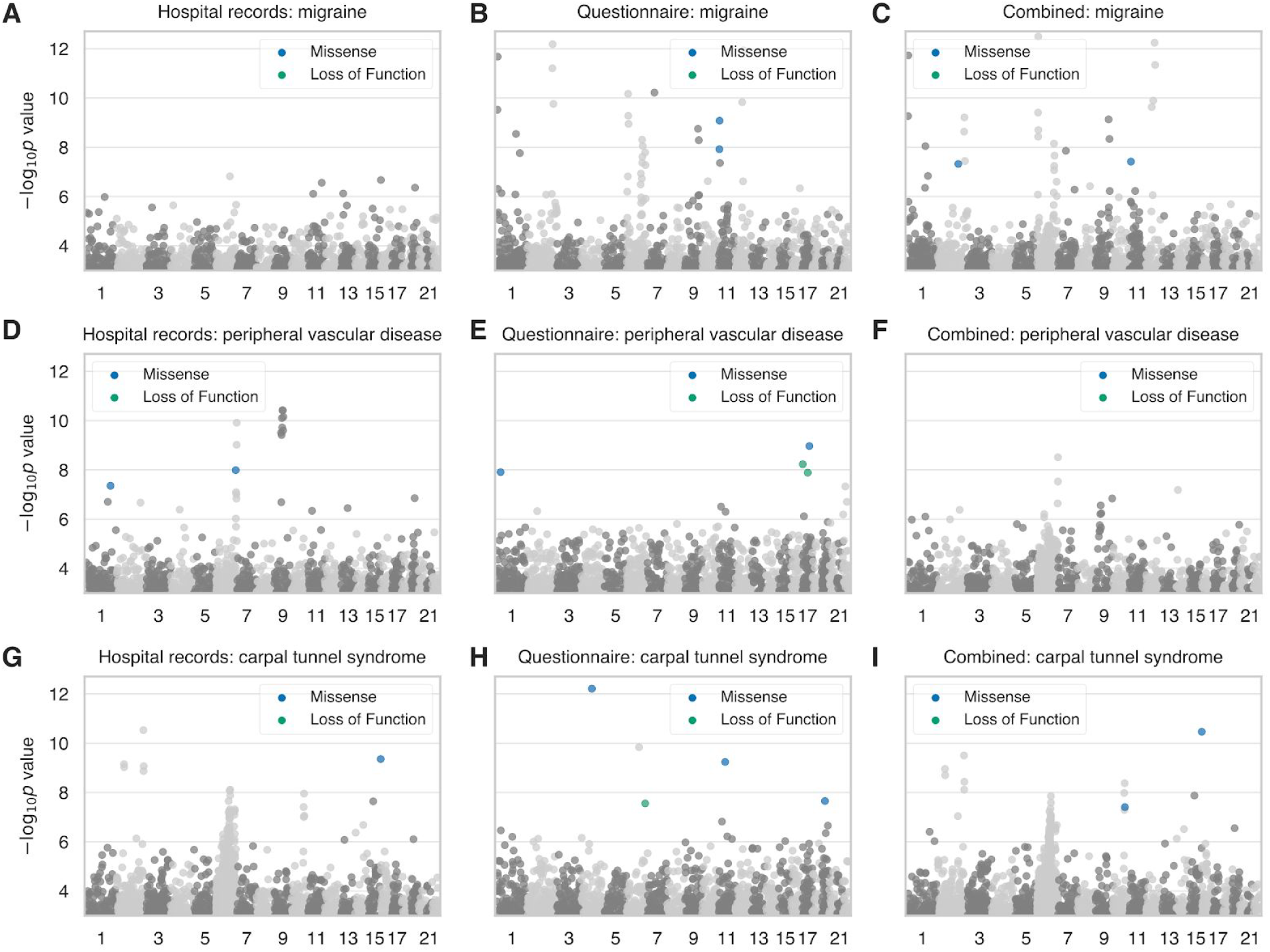
(A-C) Manhattan plots for migraine where cases were ascertained from hospital records (A), questionnaire responses (B), or both methods combined (C). (D-F) Manhattan plots for peripheral vascular disease where cases were ascertained from hospital records (D), questionnaire responses (E), or both methods combined (F). (G-I) Manhattan plots for carpal tunnel syndrome where cases were ascertained from hospital records (G), questionnaire responses (H), or both methods combined (I). For all panels, loss of function and missense variants with p<5e-8 are colored blue and green, respectively. Grey dots indicate all other variants.

**Figure S4.**
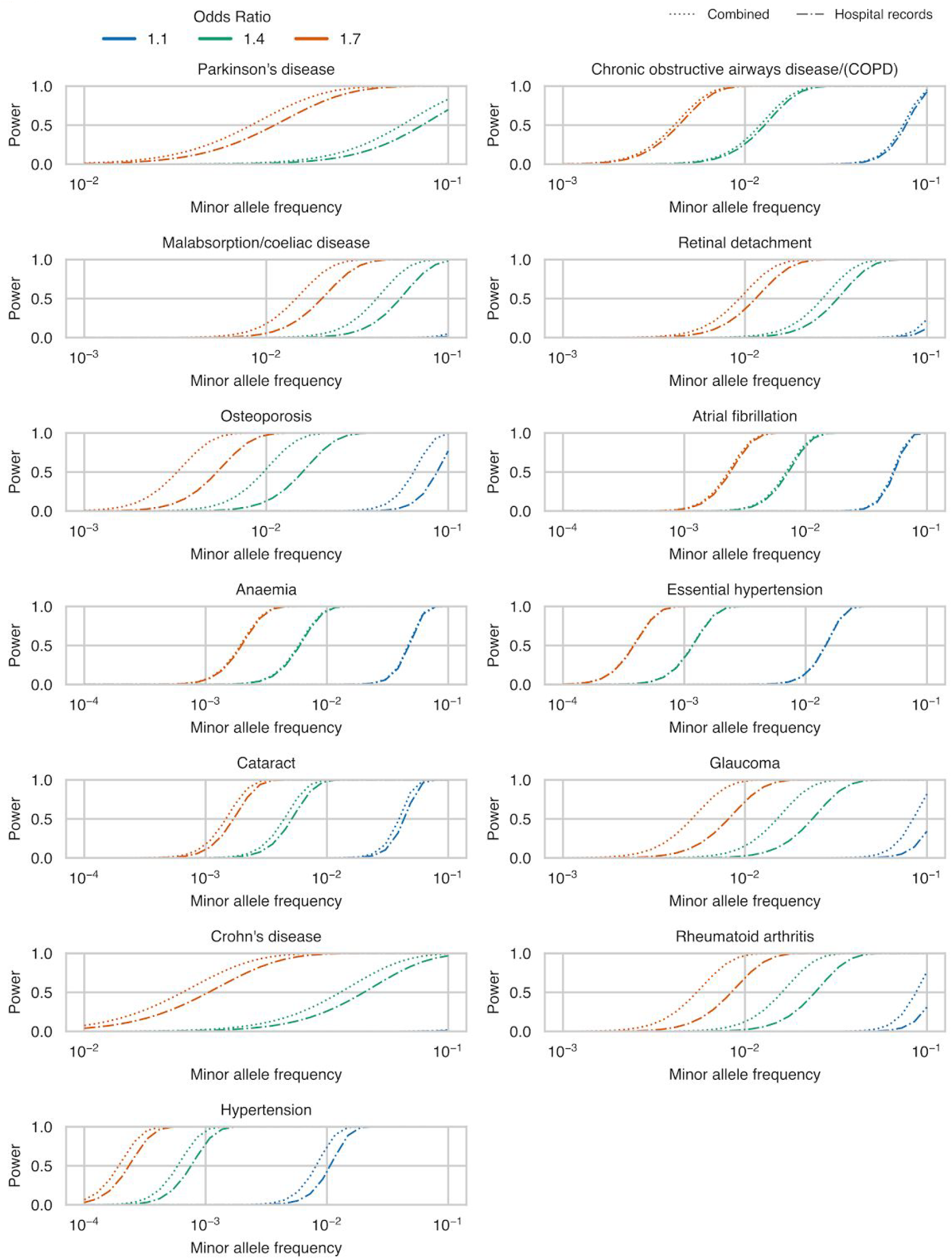
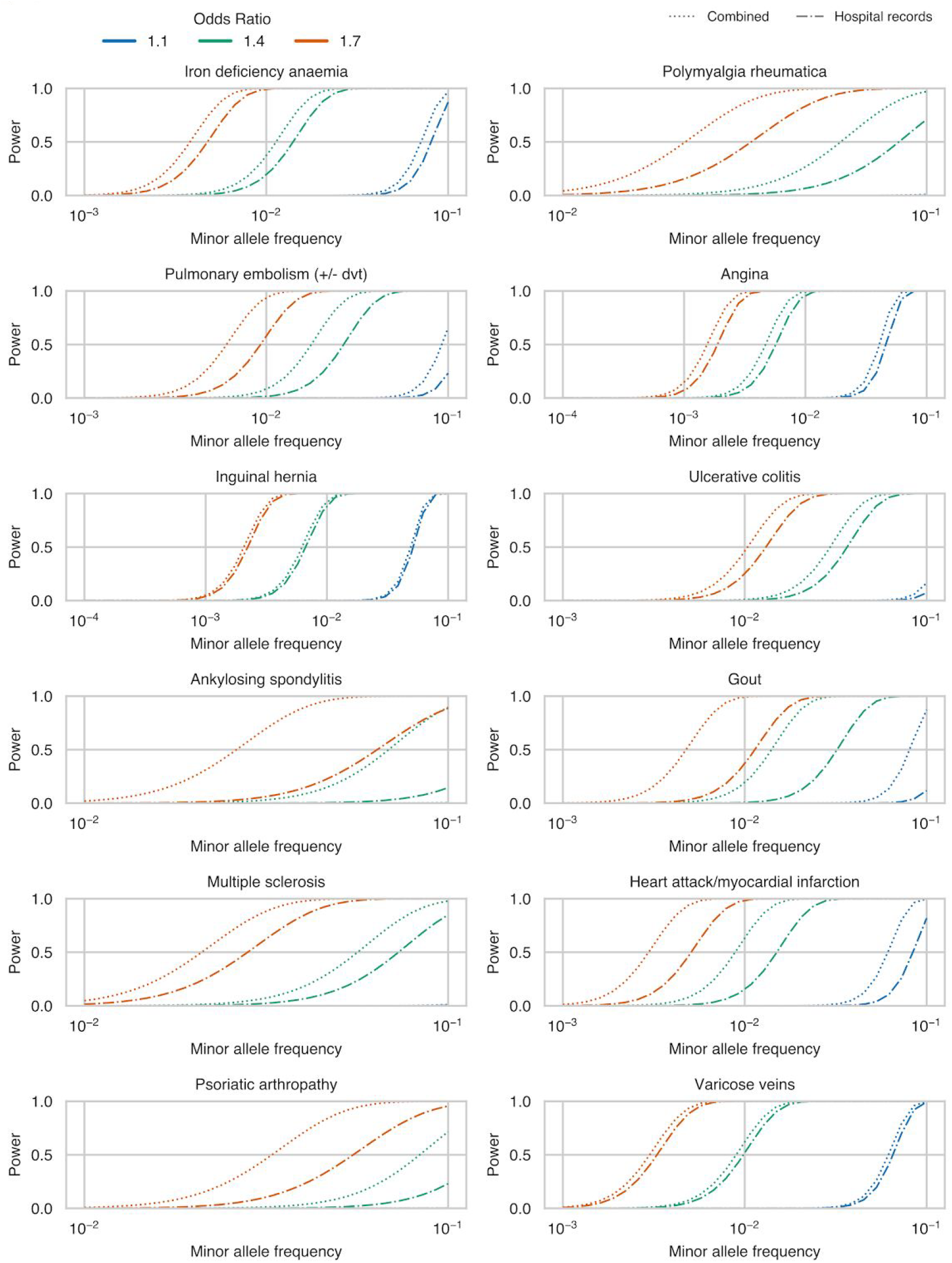
Power to detect rare risk associations among white British subjects in the UK Biobank using cases ascertained using only hospital records (dash-dot lines) or ascertained using hospital records and questionnaire responses (dotted lines). All phenotypes plotted had a mean posterior genetic correlation of at least 0.8.

**Figure S5.**
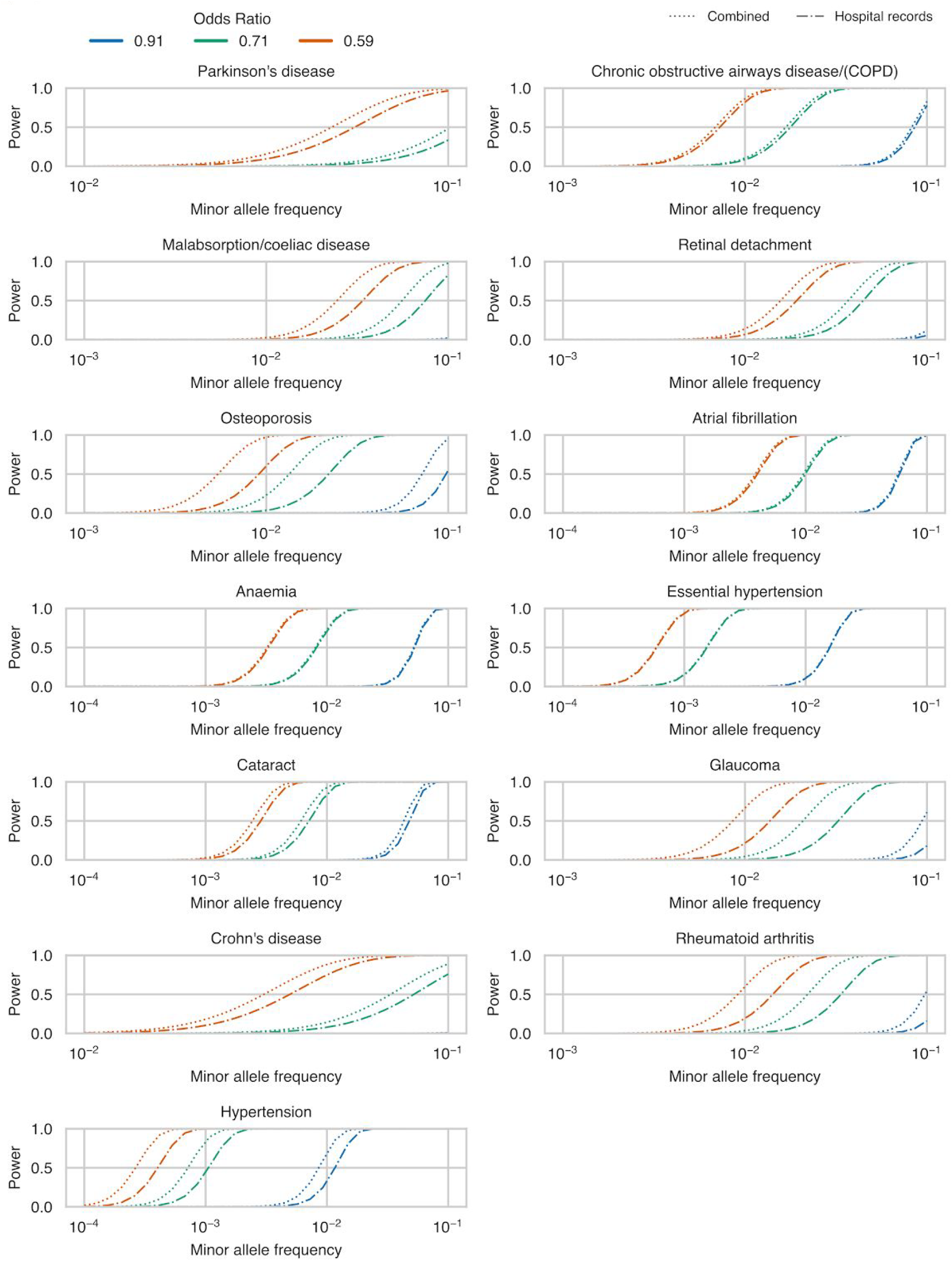
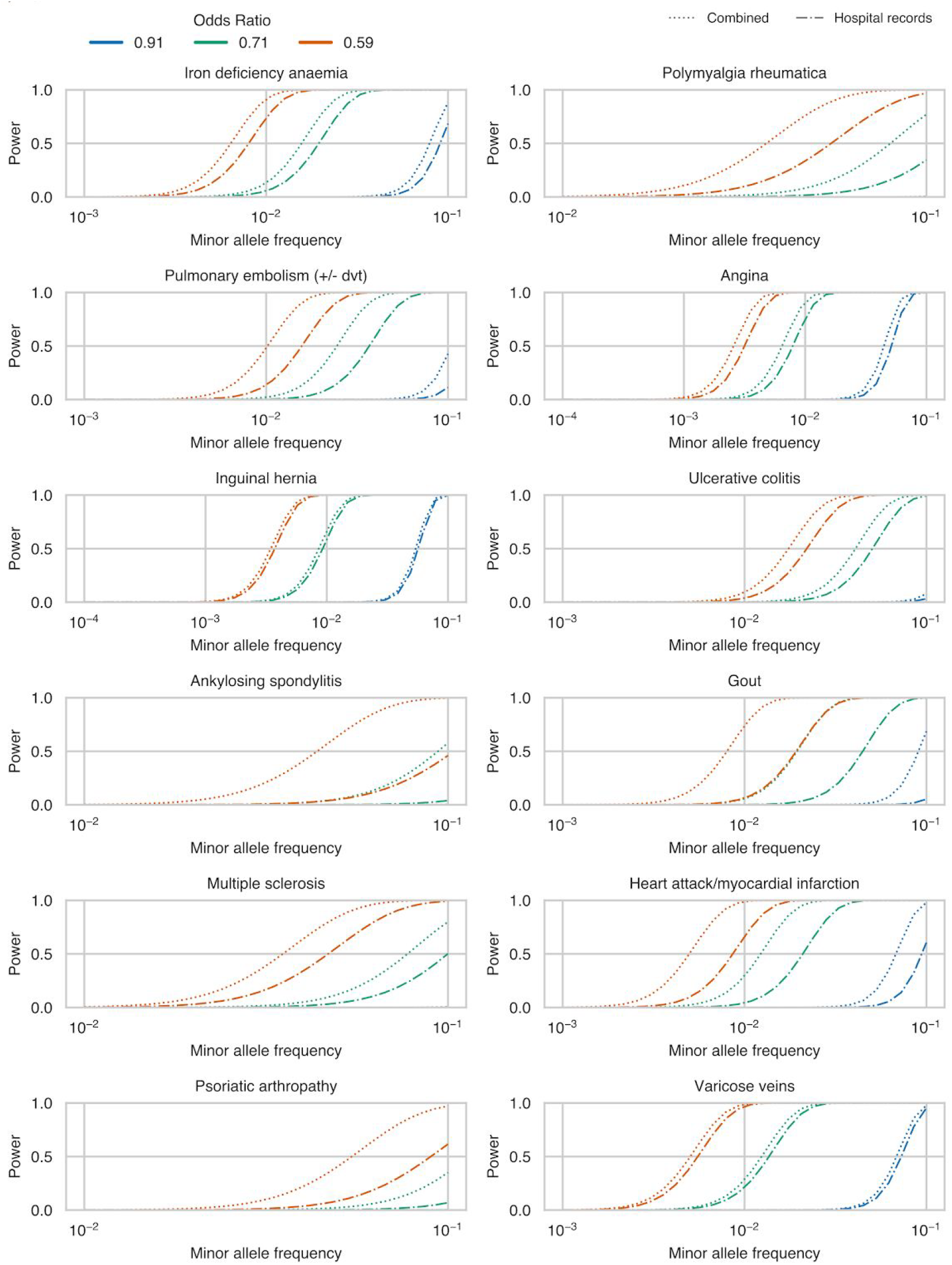
Power to detect rare protective associations among white British subjects in the UK Biobank using cases ascertained using only hospital records (dash-dot lines) or ascertained using hospital records and questionnaire responses (dotted lines). All phenotypes plotted had a mean posterior genetic correlation of at least 0.8.

**Figure S6.**
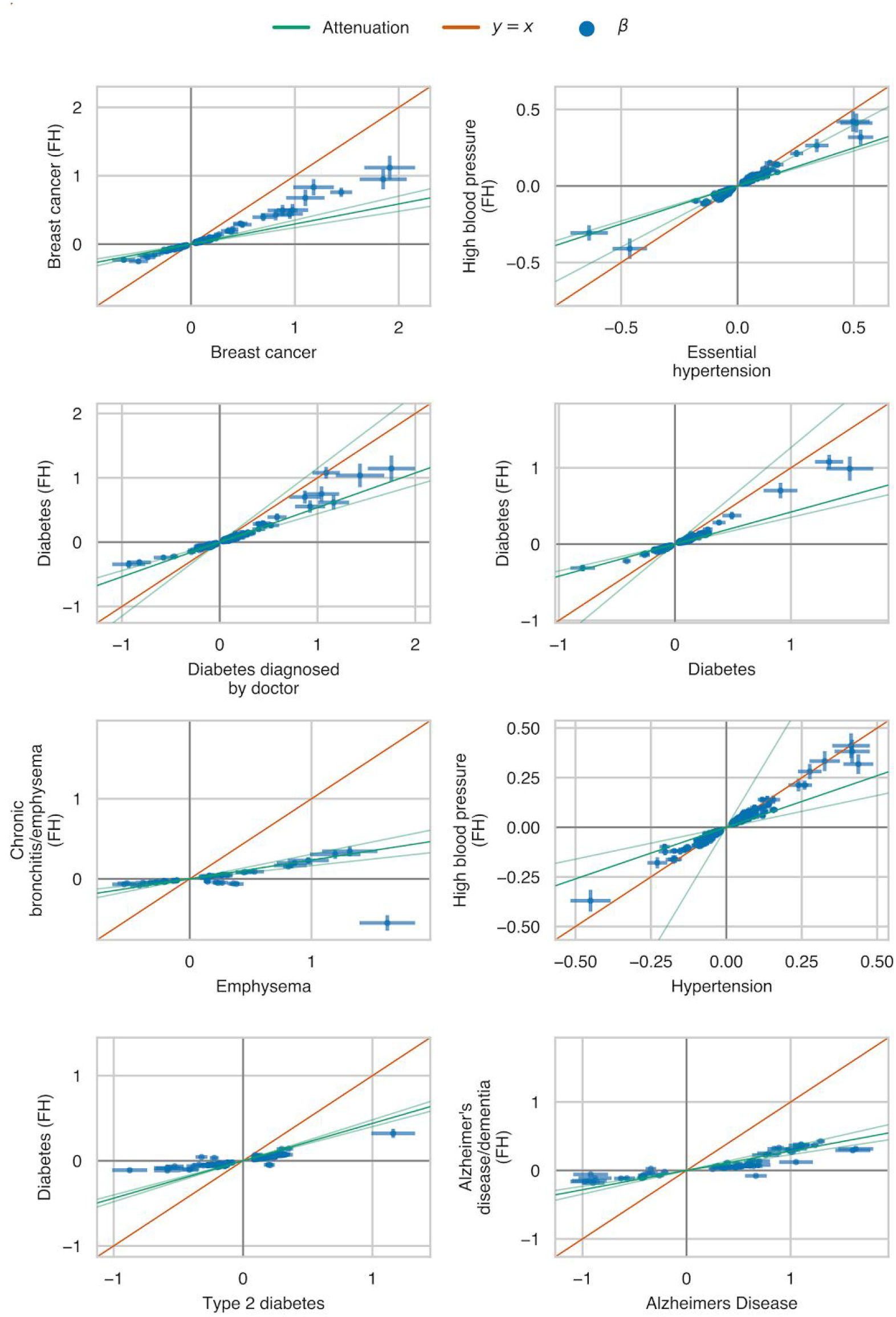
Attenuation estimates and estimated effect sizes for GWAS summary statistics from eight traits where cases were defined by either combined hospital record/verbal questionnaire data or family history of disease.

**Figure S7.**
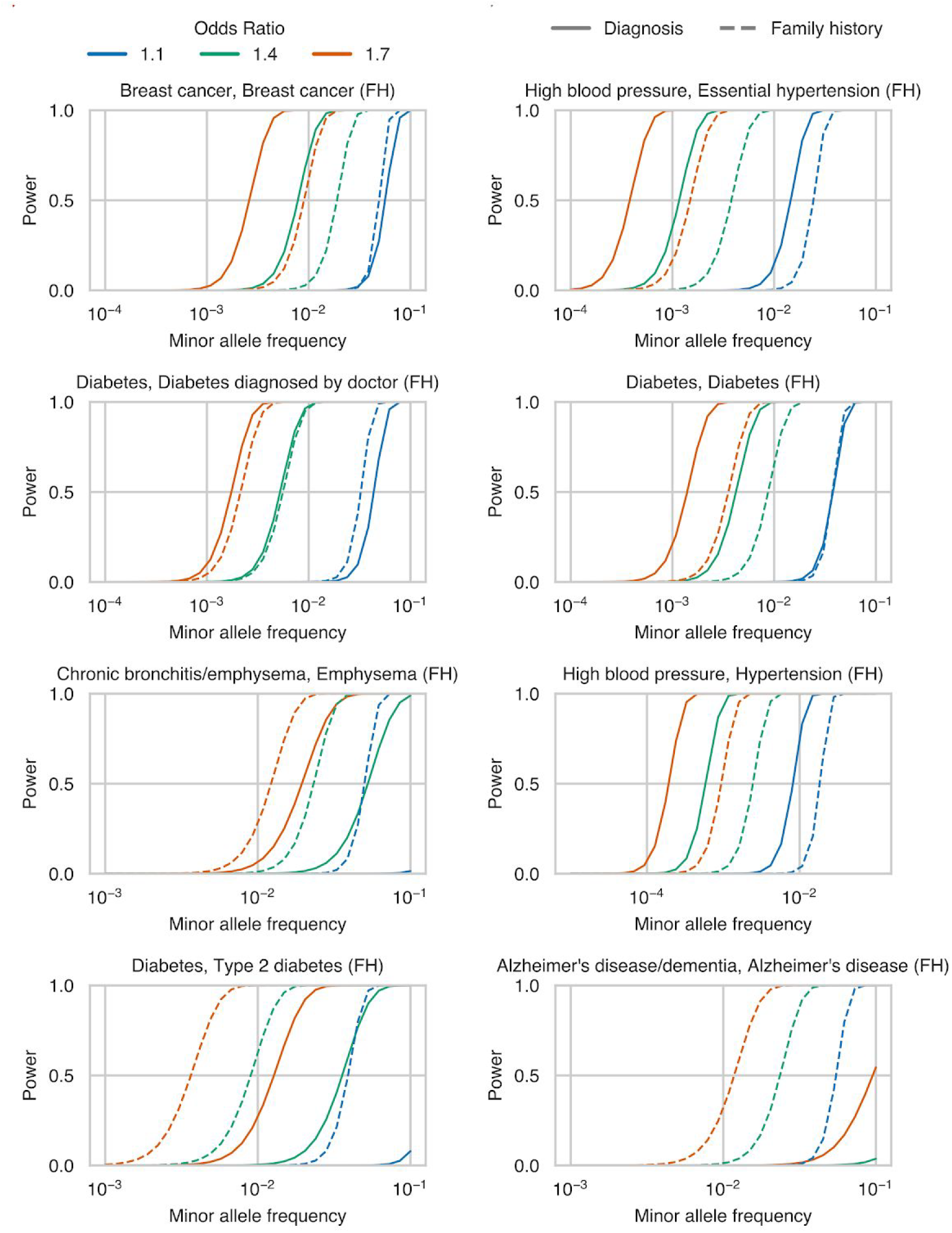
Power to detect rare risk associations among white British subjects in the UK Biobank using cases ascertained using hospital records and questionnaire responses (solid line) or family history of disease (dashed).

**Figure S8.**
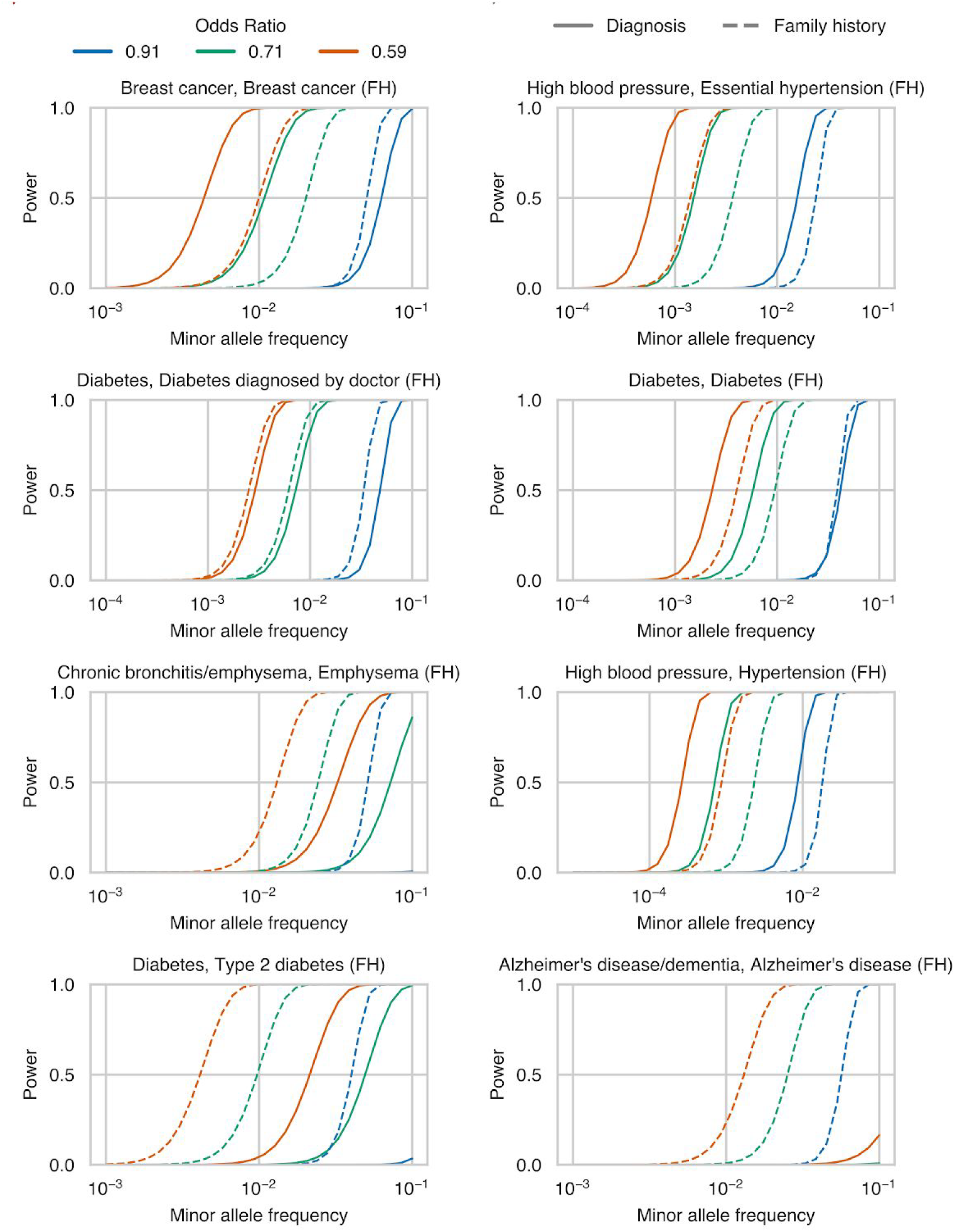
Power to detect rare protective associations among white British subjects in the UK Biobank using cases ascertained using hospital records and questionnaire responses (solid line) or family history of disease (dashed).

### Supplemental Tables

Table S1. Number of cases ascertained by hospital records, verbal questionnaire responses, and family history of disease.

Table S2. MVPMM genetic parameter estimates for comparisons of GWAS using hospital records versus questionnaire data, combined hospital records and questionnaire data versus either hospital records or questionnaire data, and family history GWAX versus combined hospital records and questionnaire data.

